# Cancer Cell’s Seven Achilles Heels: Considerations for design of anti-cancer drug combinations

**DOI:** 10.1101/2023.01.02.522527

**Authors:** Valid Gahramanov, Frederick S. Vizeacoumar, Alain Morejon Morales, Keith Bonham, Meena K. Sakharkar, Franco J. Vizeacoumar, Andrew Freywald, Michael Y. Sherman

## Abstract

Loss of function screens using shRNA and CRISPR are routinely used to identify genes that modulate responses of tumor cells to anti-cancer drugs. Here, by integrating GSEA and CMAP analyses of multiple published shRNA screens, we identified a core set of pathways that affect responses to multiple drugs with diverse mechanisms of action. This suggests that these pathways represent “weak points” or “Achilles heels”, whose mild disturbance should make cancer cells vulnerable to a variety of treatments. These “weak points” include proteasome, protein synthesis, RNA splicing, RNA synthesis, cell cycle, Akt-mTOR, and tight junction-related pathways. Therefore, inhibitors of these pathways are expected to sensitize cancer cells to a variety of drugs. This hypothesis was tested by analyzing the diversity of drugs that synergize with FDA-approved inhibitors of the proteasome, RNA synthesis, and Akt-mTOR pathways. Indeed, the quantitative evaluation indicates that inhibitors of any of these signaling pathways can synergize with a more diverse set of pharmaceuticals, compared to compounds inhibiting targets distinct from the “weak points” pathways. Our findings described here imply that inhibitors of the “weak points” pathways should be considered as primary candidates in a search for synergistic drug combinations.

## INTRODUCTION

Drug cocktails are being routinely used in anti-cancer therapies to improve tumor elimination, and reduce chances of relapse and treatment resistance. [1]. The majority of drug combinations that have been used in medical practice were selected in trial-and-error-based searches, mostly testing combinations of existing anti-cancer agents in pre-clinical models and patients [2,3]. More recently, there have been attempts to conduct massive *in vitro* screens for synergistic drug combinations [4–6]. Unfortunately, such screens often have limited success, because of insufficient throughput. In addition, investigators have been approaching the problem in a more rational way, by predicting synergistic combinations based on the established mechanisms of action of anticancer drugs. For example, in case of insufficient effects of ERBB2 inhibition due to the compensatory overexpression of the EGF receptor (EGFR) in cancer cells, administration of the ERBB2 inhibitor, Herceptin, is often combined with suppressing EGFR activity [7–10]. Beyond all these strategies, sophisticated computational analyses of treatment-induced transcriptome modifications and machine learning algorithms have been applied to find novel potent drug combinations [5,6]. So far, these efforts have not produced effective drug combinations for most cancer types, especially for advanced and metastatic tumors. Therefore, new and more effective research approaches have to be developed.

Lately, shRNA and CRISPR screens have been employed to comprehensively characterize cellular effects of newly developed and more traditional therapeutic compounds, and to identify genes associated with resistance to their cancer-suppressing action. Such screens have a strong potential to facilitate search for novel drug combinations [11–13]. In our earlier work, people used shRNA screens to identify genes involved in responses to a novel Hsp70 inhibitor JG-98 [14] and to an antibiotic gentamycin [15]. Surprisingly, a set of common genes was identified in these studies, although the screens were focused on two chemicals that were entirely different both structurally and in their mechanisms of action. Based on this unexpected finding, we suggested that there may be a core set of genes that constitute “weak points” of the cell, and depletion of such genes may make cells vulnerable to drugs and stressors with different mechanisms of action. Knowledge of these “weak points” could be very helpful in designing novel drug cocktails. Here, we addressed the “weak points” hypothesis by analyzing multiple available datasets from cancer-related shRNA screens and previously tested drug combinations.

## RESULTS

### Global view of screens and enrichment analyses

To determine if certain genes or gene sets represent common “weak points” of cancer cells that could be targeted to enhance effectiveness of existing therapeutic compounds, we pooled hits from shRNA screens done against 8 unrelated drugs (Supplementary Table S1) [14,16–18].

These data were subjected to the Gene Set Enrichment Analysis (GSEA) to identify pathways that affect sensitivity to the drugs used in the screens. To assess correlation between sizes of gene sets present in the shRNA libraries and the numbers of the identified hits belonging to the established pathways, we employed the correlation analysis. A linear correlation between the number of genes present in shRNA libraries and the number of hits in the screens is shown in Fig.1A. The only outlier was a screen with the Top1 inhibitor irinotecan that showed significantly higher fraction of pathways enriched genes, compared to other screens.

**Figure 1.**
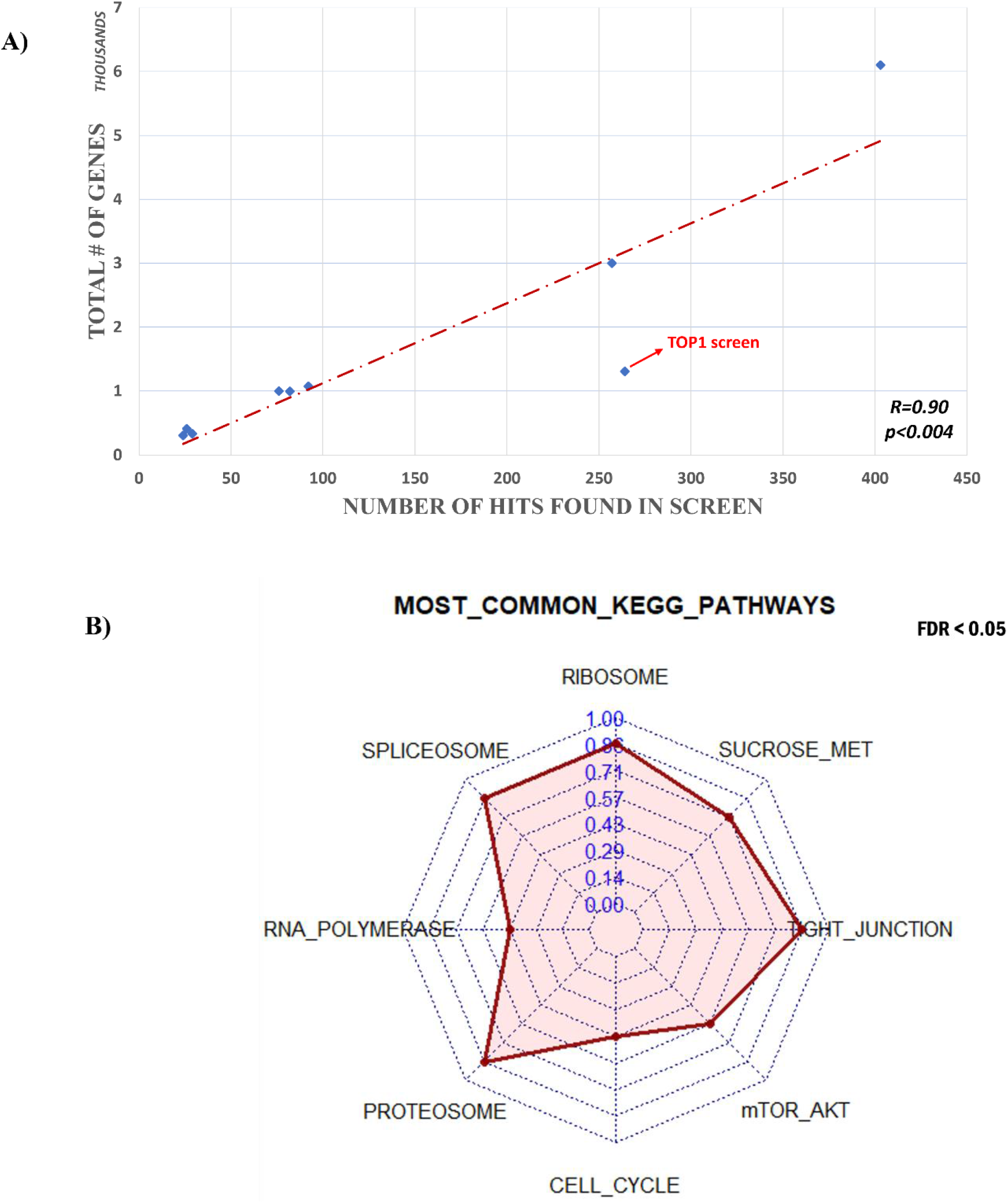
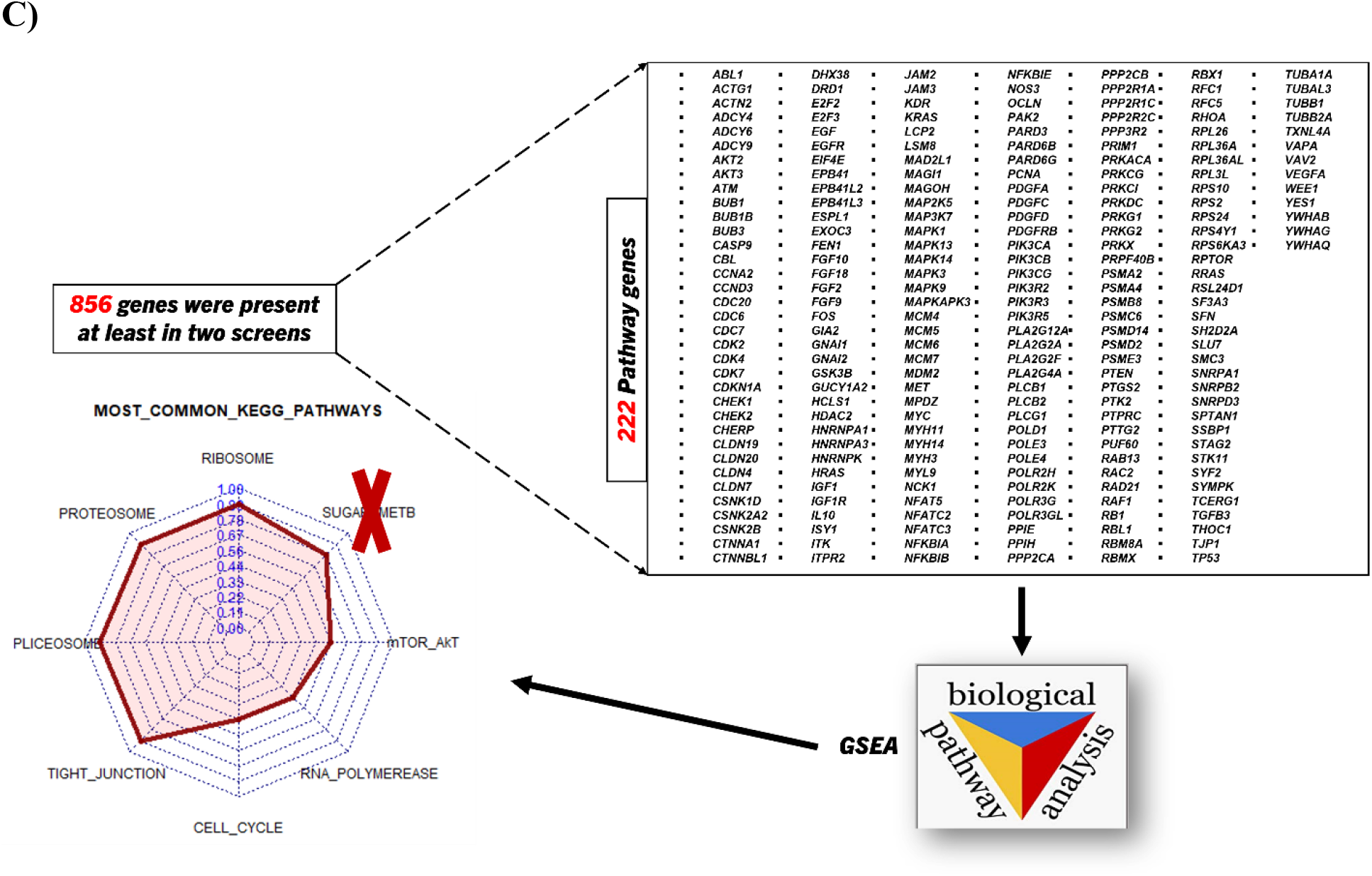
**A) Global view of shRNA screening data analysis.** Represents the linear regression of 8 shRNA screening library correlation with the found hits. Bigger library has more probability to have higher number of hits. **B) GSEA analysis of shRNA screens.** Radar plot describes the signaling pathways of sensitized genes of shRNA screens. The density of red color indicates the commonality of pathways in different screen. Dotted line represents individual shRNA screens. **C) Overall representation of pathway analysis of overlapped genes among shRNA screens**.

### Identification of “weak points”pathways

To find biological processes that became crucial upon administration of various treatments, we compared lists of pathways, whose inhibition sensitized cells in different screens. This approach revealed pathways common in at least 3, at least 4 and at least 5 screens (Table S2). Notably, in the analysis of these pathways, one needs to consider that their designation in GSEA is somewhat arbitrary. In other words, same genes could belong to different pathways defined in GSEA. Indeed, analysis of gene sets that constitute pathways indicated in Fig. 1B revealed that the designation of T-cell signaling, VEGF signaling, mTOR, cytosolic DNA sensing, pathogenic E. coli infection and Gap-junction pathways was due to the presence of the same genes found in shRNA screens. Therefore, we considered these pathways together as the Akt-mTOR pathway. Overall, we found several core pathways, which when suppressed, increase sensitivities of cancer cells to drugs in at least three independent screens. This list included ribosome, spliceosome, transcriptomic, proteasome, cell cycle, Akt-mTOR, starch and sucrose metabolism, and tight junction pathways. Among pathways common in at least four screens, we lost the transcription pathway, and pathways common in at least five screens also did not include the mTOR pathway (Fig. 1B, Table S2.)

To complement this work, we performed a different analysis by evaluating the overlapping genes identified in the shRNA screens. Despite the high diversity of tested drugs in the screens, there were 856 genes present in at least 2 screens. Considering that screens often produce false-negative and false-positive hits, we enhanced the stringency of our analysis by concentrating only on the overlapping genes that belong to common pathways. This approach allowed us to narrow our search to 222 candidates (Fig. 1C).

By analyzing 222 recurrently found genes using the Molecular Signature Database [19], we identified common pathways that appeared to be responsible for the resistance to several drugs Overall, they proved to be similar to common pathways identified via the direct comparison of pathways in shRNA screens. As discussed above, we considered T-cell signaling, VEGF signaling, mTOR and Gap-junction pathways collectively, as the Akt-mTOR pathway. Similarly, we considered DNA replication as part of the Cell Cycle pathway. Taken together, our two independent strategies pointed towards seven core pathways including ribosome, spliceosome, transcription, proteasome, cell cycle, Akt-mTOR and tight junction, inhibition of which sensitized cancer cells to various drugs. Overall, these data indicate that suppression of certain specific pathways in cancer cells may make them more sensitive to drugs with distinct mechanisms of action and therefore, these pathways may represent “weak points” of cancer cells.

### Inhibitors of the “weak points” pathways synergize with more diverse sets of drugs compared to inhibitors of other pathways

The hypothesis of “weak points” in the cell makes a clear prediction - drugs that synergize with inhibitors of the “weak points” pathways should be more diverse compared to sets of drugs that synergize with therapeutics that affect targets other than the “weak points”. To test this prediction, we collected published data on the drugs that synergize with “weak point” pathway inhibitors (test therapeutics), and with control therapeutics that target other pathways (control therapeutics). Certain possible biases should be considered in such metadata analysis, including (a) investigator biases related to preexistent information about drug effects in choosing combinations to test, i.e., non-random choice of tested drug combinations, and (b) bias related to the novelty of tested therapeutics, which results in only a few reported synergistic combinations (Fig. 2A). We attempted to overcome both types of biases by choosing for our analysis only therapeutics that had more than 30 published papers that discussed different established synergistic combinations. Based on this, we excluded translation and splicing pathways, since their targeting is new in the clinical practice and there is no sufficient number of reported synergistic drugs. The tight junction pathway was excluded because there are no known drugs that target this pathway. Therefore, we selected inhibitors of proteasome (bortezomib), transcription (CDK9 inhibitor), cell cycle (abemaciclib) and Akt-mTOR (rapamycin) pathways as test therapeutics for the metadata analysis. Drugs that synergize with inhibitors of these pathways are shown in Supplementary Table S3. As control therapeutics that inhibit targets other than the weak points, we chose inhibitors of topoisomerase I (irinotecan), EGFR (erlotinib), genotoxic agent (cisplatin) and antimetabolite (fluorouracil).

**Figure 2.**
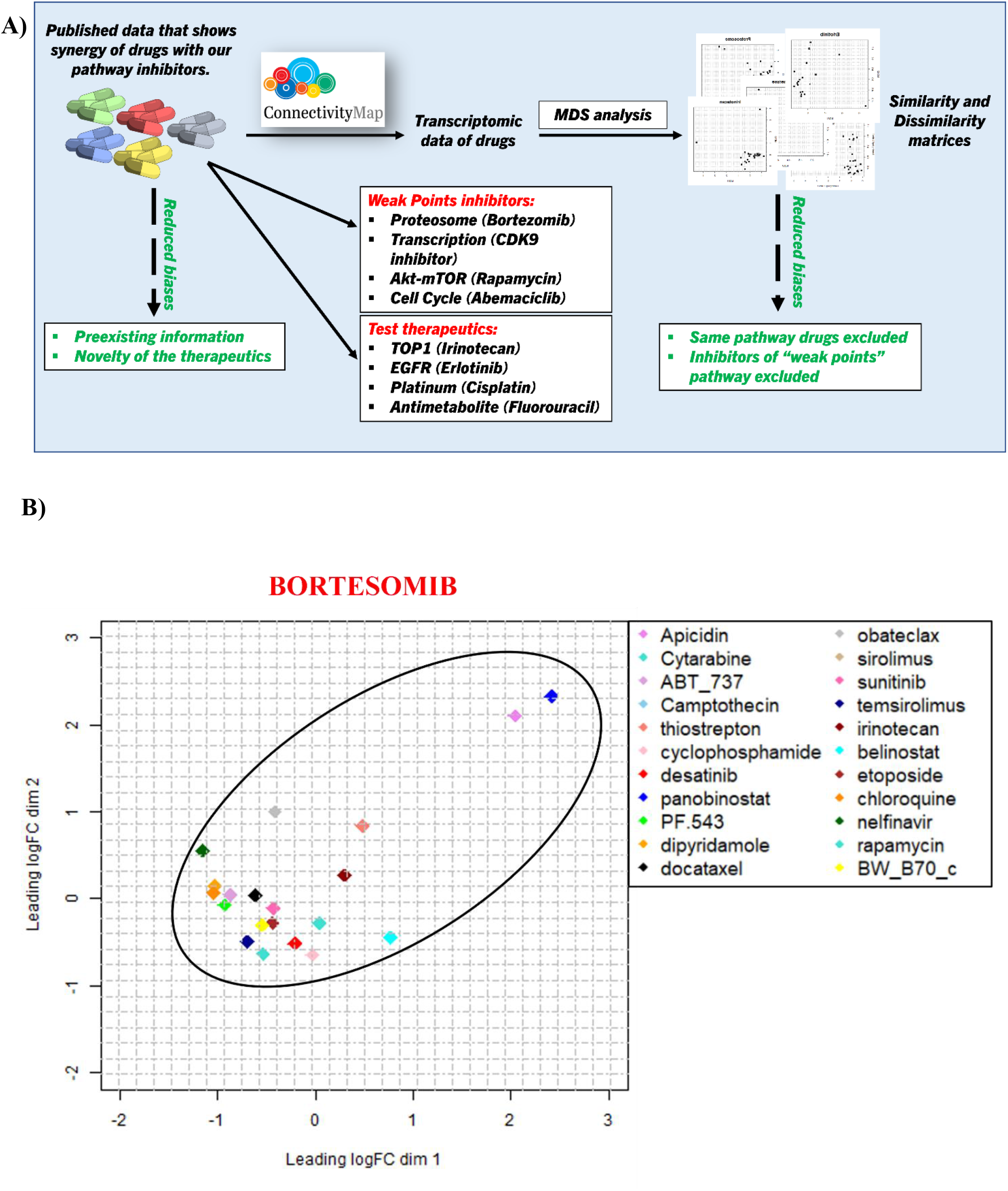
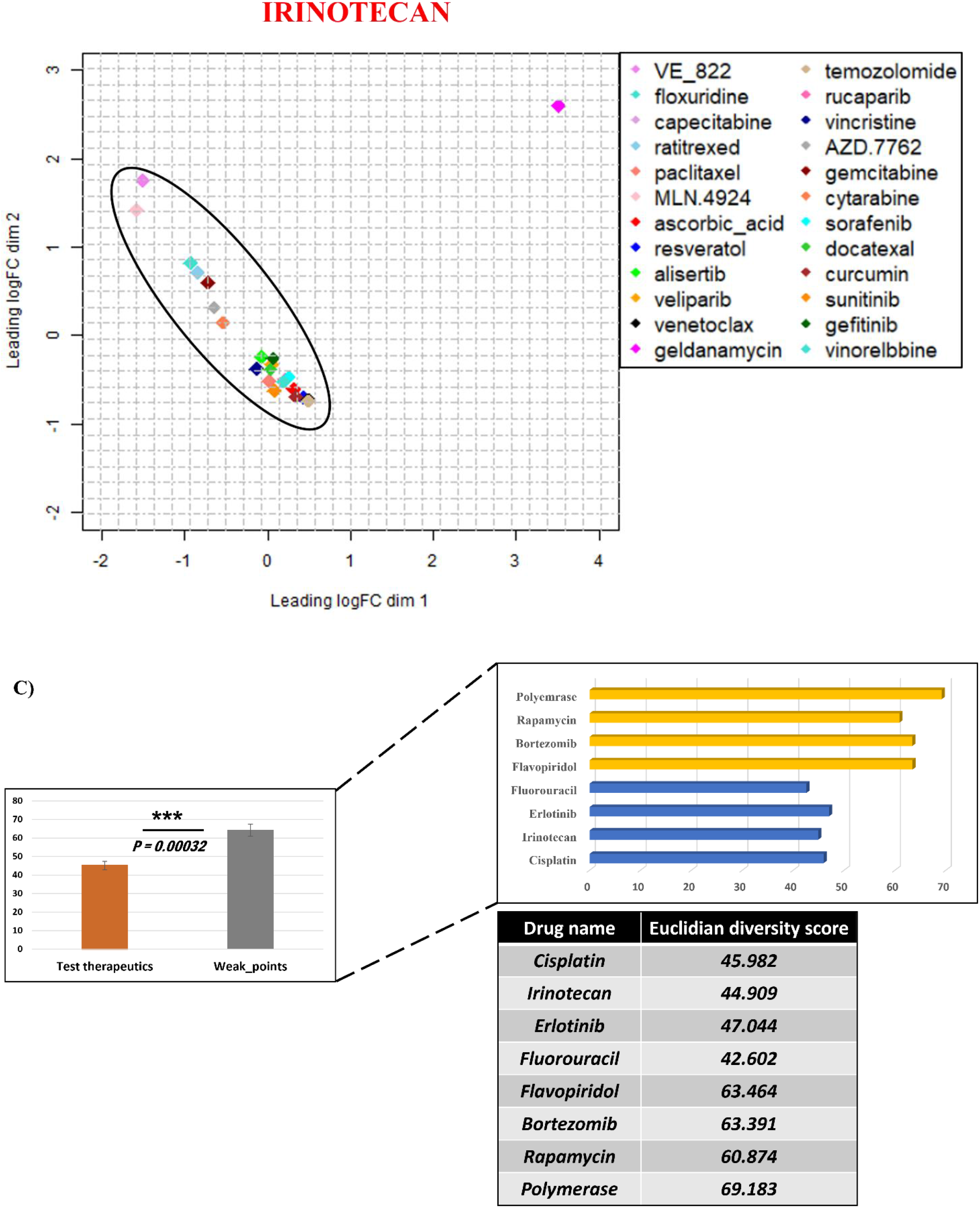
Finding synergistic drugs. **A)** Graphic represents overall summary of the stages to find out synergistic drugs based on defined “weak point” pathways. **B)** MDS plot describes the diversity of drugs that synergizes with so-called “weak point” pathways inhibitor on the left and “test therapeutics” (control). **C)** Bar plots represents all the statistic transcriptional profile of drugs that synergizes both “weak point pathways” inhibitor and “test therapeutics”. * - *p-value<0.05* Statistical analysis has been done via Student’s t Test by considering 4 drugs per each group (n=4).

We decided to approach the diversity based on the transcriptomics effects of the synergistic drugs. Accordingly, we derived the transcriptomic effects of drugs listed in the Supplementary Table S3 from the Connectivity Map project (CMAP) [20]. Since in this database, transcriptomics effects of drugs are compared in multiple cancer lines, we standardized the analysis by choosing effects of all drugs of interest on a single breast cancer cell line, MCF7. Of note, these drugs showed very similar transcriptomic effects in other cancer cell lines. Upon extraction of the transcriptomics information, we performed multidimensional scaling analysis to assess similarities and dissimilarities of transcriptomic profiles of drugs that synergize with test therapeutics targeting the “weak points” and control pathways. To statistically estimate the diversity of the listed drugs that synergize with test therapeutics we used Euclidean distance in MDS plot to score average distances between synergizing drug effects, in other words, the degree of their clustering in a 2-D graph. In the MDS analysis, we excluded (a) drugs that affect the same pathways as test therapeutics, and (b) inhibitors of the other “weak point” pathways, since they would also increase the diversity through their action (Fig. 2A).

As expected, the transcriptional diversity of drugs synergized with inhibitors of the “weak points” pathways was higher compared to the diversity of drugs that synergize with control therapeutics, and this difference was highly statistically significant (Fig. 2B, 2C, Supplementary Figure S1). These findings support the hypothesis that at least proteasome, transcription and Akt-mTOR pathways are Achilles points of the cell, partial inhibition of which makes cancer cells more sensitive to a variety of drugs.

## Conclusion

Here, we uncover that certain pathways represent “weak points” of cancer cells, mild disturbance of which is likely to make cells more vulnerable to dysregulation of homeostasis by a variety of stressful conditions and drugs. This finding provides a valuable information that should assist in searches for optimal synergistic cancer drug combinations.

## Discussion

Combination therapies are viewed as most effective strategies of treating cancer patients [21–24]. Here, based on multistep bioinformatics analysis of previously published and unpublished pooled shRNA screens, we determined that partial depletion of genes belonging to a limited set of pathways enhances sensitivities of cancer cells to various therapeutic compounds. Surprisingly, these drugs were unrelated in their mechanisms of action, suggesting that partial suppression of the identified pathways make cells more sensitive to general disturbances of their homeostasis. Therefore, we denominate them “weak points” of the cell.

Analysis of genes belonging to the “weak points” pathways uncovered a relation to “cancer essentiality genes” [25–27]. Indeed, previous study used a large-scaled shRNA screen to identify genes essential for the survival of cancer cells. It uncovered 297 of such genes (so-called cancer essentiality genes) crucial for the survival and proliferation of cancer cells in 72 cell lines, but not non-malignant cells [25]. While some of these genes have important functions in a number of cellular responses, their partial suppression by shRNA-based silencing proved to be selectively lethal for cancer cells [28]. We observed a strong overlap between cancer essentiality pathways and pathways that affect sensitivity to various drugs in our analysis of shRNA screens, including ribosome, splicing, transcriptome, cell cycle and proteasome. It is likely, that in the publication identifying the essentiality genes relatively strong gene silencing was achieved that was sufficient for the loss of viability of cancer cells. In contrast in the published drug-related screens that we used in our study a milder gene downregulation must have been applied, which allowed cancer cells to live, while making them more sensitive to certain compounds. Interestingly, two “weak points” pathways, including Akt-mTOR and tight junction identified in our work, were not found among the cancer essentiality pathways.

The concept of “weak points”, i.e., selected pathways that can be suppressed to sensitize cancer cells to distinct treatments predicts that inhibitors of these pathways should synergize with highly diverse set of drugs. To address this question, we examined, how diverse synergistic drugs to inhibitors of the “weak points” pathways and inhibitors of other pathways are. In other words, we quantitatively evaluated the diversity of drugs synergistic to these therapies. It is unclear, how to do such evaluation based on mechanisms of the drugs action. For example, it is unclear how to estimate whether the pair of an inhibitor of proteasome and EGFR is more or less diverse than the pair of inhibitors of mTOR and topoisomerase I. We approached the diversity based on the transcriptomics effects of the synergistic drugs. Extraction of transcriptomics effects of selected drugs from the CMAP database indeed allowed to compare and score diversity levels. We realize that functional comparison would probably be more adequate, but as noted above, such a comparison does not allow scoring. Our data also support the idea that transcriptomic profiles of anti-cancer compounds can be used as an effective tool in the search of effective drug combinations.

More specifically, our work strongly suggests that the identified “weak points” pathways should be given a priority as candidates for developing new combination therapies.

## MATERIALS AND METHODS

### Data resource and study design

Data on shRNA screens of drug effects were collected from published works. Genes depletion which sensitizes to drugs in each screen is shown in Supplementary Table S1.

### Gene Set Enrichment Analysis (GSEA) and Connectivity Map (CMAP)

The entire list of sensitizing genes of screening was ranked according to fold change and used as input to Gene Set Enrichment Analysis (GSEA) [29,30]. GSEA was employed against both the hallmark gene-set signature and curated gene sets. The hallmark and the curated gene sets were obtained from the Molecular Signature Database v7.2 (http://www.gsea-msigdb.org/gsea/msigdb/index.jsp). We have taken into consideration gene sets with a p-value < 0.05 and an FDR cutoff of 25% in most of the screening data, which is appropriate in GSEA analysis due to the relatively small number of gene sets being analyzed (50) [29]. In some screens, the FDR cutoff was increased to 35-40% due to the least number of hits found in the original data. shRNA expression data were collected from GEO Datasets databases [31]. Transcriptomic data of drug effects are collected from Connectivity Map databases [32,33]. To make all the transcriptomic profile analyses unbiased we extracted the transcriptomic effects of each drug after 24h treatment in the MCF7 cell line.

### Drug synergy analysis

PubMed datasets [34] were selected for identifying synergistic drug combinations. We use keywords, e.g., ‘Proteasome inhibitor’ AND ‘synergy in cancer’ to find publications that contain information about effects of drugs synergistic with proteasome inhibitors. Further, we followed the same searching method for the rest pathway inhibitors - ‘CDK9’, ‘Cisplatin’, ‘Irinotecan’, ‘Rapamycin’, ‘Cell cycle’.

### Multidimensional Scaling

To check the similarity and dissimilarity of transcriptomic effects of drugs, we utilized multidimensional scaling analysis [35] with *edgeR* [36] and *limma* [37] packages in R programming language. Distances of the drugs’ transcriptomic data on dimensional plot were calculated based on the Euclidean distance algorithm [38,39].

### Statistical Analysis

Statistical analysis was performed in R. For the calculation of Euclidian Distance, we used Pythagorean equation – 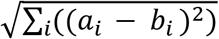. For observations a and b measured on many dimensions. Student’s t test was used to evaluate the statistical significance of difference in two groups. P-value less than 0.05 was considered as significant. Statistical differences are representative of consideration of three observations per groups (n=3).

## Author Contributions

The metadata analysis was designed and performed by **V.G**. Unpublished shRNA screens was performed by **A.M.M., K.B., M.K.S., S.K**. Data handling and analysis, signaling pathway analysis was performed by **V.G.** Valuable suggestion in data analysis were provided by **F.J.V., F.S.V., M.S.D.** Manuscript was prepared by **V.G., M.S., A.F.** Manuscript reviewing and editing done by **M.S., A.F., F.J.V.** Manuscript was approved by all co-authors. All authors have read and agreed to the published version of the manuscript.

## Data Availability Statement

The manuscript contains supplementary files and has been provided separately. Raw and processed data are in process of storing in GEO database website and will be publish during the reviewing process of the original manuscript. Codes and pipelines will be made upon available upon request.

## Abbreviations

GSEA: Gene Set Enrichment Analysis
CMAP: Connectivity Map
CRISPR: Clustered Regularly Interspaced Short Palindromic Repeats
RNA: Ribonucleic acid
DNA: Deoxyribonucleic acid
shRNA: short hairpin RNA
MDS: Multidimensional Scaling
FDR: False Discovery Rate
GEO: Gene Expression Omnibus
VEGF: Vascular endothelial growth factor
EGF: epidermal growth factor
EGFR: epidermal growth factor receptor.

