## Supplementary material for "Cancer Cell’s Seven Achilles Heels: Considerations for design of anti-cancer drug combinations": SUPPLEMENTARY FIGURES AND TABLES.docx

**Table S2: Identification of common signaling pathways among the shRNA screening data.** The tables below represent the pathways that were found in at least 2, 3, 4, and 5 independent screens.

Pathways are present at least in 2 shRNA screens.

KEGG_TGF_BETA_SIGNALING_PATHWAY

KEGG_PROPANOATE_METABOLISM

KEGG_PROTEIN_EXPORT

KEGG_TOLL_LIKE_RECEPTOR_SIGNALING_PATHWAY

KEGG_GLYCOSYLPHOSPHATIDYLINOSITOL_GPI_ANCHOR_BIOSYNTHESIS

KEGG_EPITHELIAL_CELL_SIGNALING_IN_HELICOBACTER_PYLORI_INFECTION

KEGG_LEISHMANIA_INFECTION

KEGG_PYRUVATE_METABOLISM

KEGG_TYPE_II_DIABETES_MELLITUS

KEGG_INSULIN_SIGNALING_PATHWAY

KEGG_NEUROTROPHIN_SIGNALING_PATHWAY

KEGG_PENTOSE_AND_GLUCURONATE_INTERCONVERSIONS

KEGG_DNA_REPLICATION

KEGG_HOMOLOGOUS_RECOMBINATION

KEGG_PYRIMIDINE_METABOLISM

KEGG_PHENYLALANINE_METABOLISM

KEGG_OXIDATIVE_PHOSPHORYLATION

KEGG_HUNTINGTONS_DISEASE

KEGG_ECM_RECEPTOR_INTERACTION

KEGG_MAPK_SIGNALING_PATHWAY

KEGG_ARGININE_AND_PROLINE_METABOLISM

KEGG_CYTOKINE_CYTOKINE_RECEPTOR_INTERACTION

KEGG_ABC_TRANSPORTERS

KEGG_NOD_LIKE_RECEPTOR_SIGNALING_PATHWAY

Pathways are present at least in 3 shRNA screens.

KEGG_RIBOSOME

KEGG_GAP_JUNCTION

KEGG_TIGHT_JUNCTION

KEGG_PROTEASOME

KEGG_SPLICEOSOME

KEGG_CELL_CYCLE

KEGG_VEGF_SIGNALING_PATHWAY

KEGG_T_CELL_RECEPTOR_SIGNALING_PATHWAY

KEGG_UBIQUITIN_MEDIATED_PROTEOLYSIS

KEGG_ANTIGEN_PROCESSING_AND_PRESENTATION

KEGG_BETA_ALANINE_METABOLISM

KEGG_ALANINE_ASPARTATE_AND_GLUTAMATE_METABOLISM

KEGG_ENDOCYTOSIS

KEGG_CYTOSOLIC_DNA_SENSING_PATHWAY

KEGG_PATHOGENIC_ESCHERICHIA_COLI_INFECTION

KEGG_RNA_POLYMERASE

KEGG_STARCH_AND_SUCROSE_METABOLISM

KEGG_MTOR_SIGNALING_PATHWAY

KEGG_MISMATCH_REPAIR

KEGG_NUCLEOTIDE_EXCISION_REPAIR

Pathways are present at least in 4 shRNA screens.

KEGG_RIBOSOME

KEGG_GAP_JUNCTION

KEGG_TIGHT_JUNCTION

KEGG_PROTEASOME

KEGG_SPLICEOSOME

KEGG_VEGF_SIGNALING_PATHWAY

KEGG_ANTIGEN_PROCESSING_AND_PRESENTATION

KEGG_CYTOSOLIC_DNA_SENSING_PATHWAY

KEGG_PATHOGENIC_ESCHERICHIA_COLI_INFECTION

KEGG_STARCH_AND_SUCROSE_METABOLISM

KEGG_MTOR_SIGNALING_PATHWAY

Pathways are present at least in 5 shRNA screens.

KEGG_RIBOSOME

KEGG_RIBOSOME

KEGG_GAP_JUNCTION

KEGG_TIGHT_JUNCTION

KEGG_PROTEASOME

KEGG_SPLICEOSOME

KEGG_CYTOSOLIC_DNA_SENSING_PATHWAY

KEGG_PATHOGENIC_ESCHERICHIA_COLI_INFECTION

KEGG_STARCH_AND_SUCROSE_METABOLISM

**Figure S1:** **B)** MDS plots describe the diversity of drugs that synergizes with so-called “weak point” pathways inhibitors – Cisplatin, Erlotinib, and “test therapeutics” (control) – Polymerase inhibitor and Rapamycin.

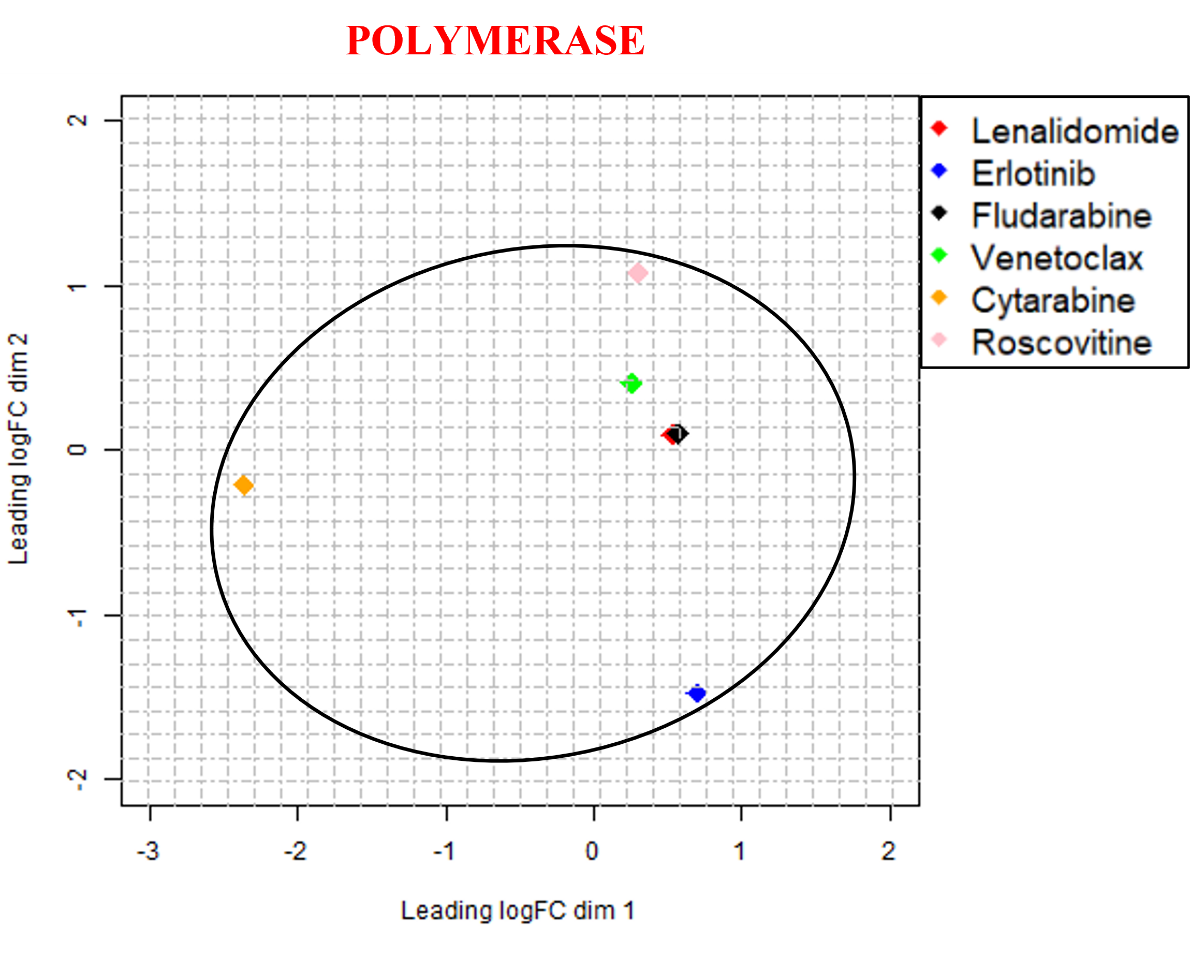

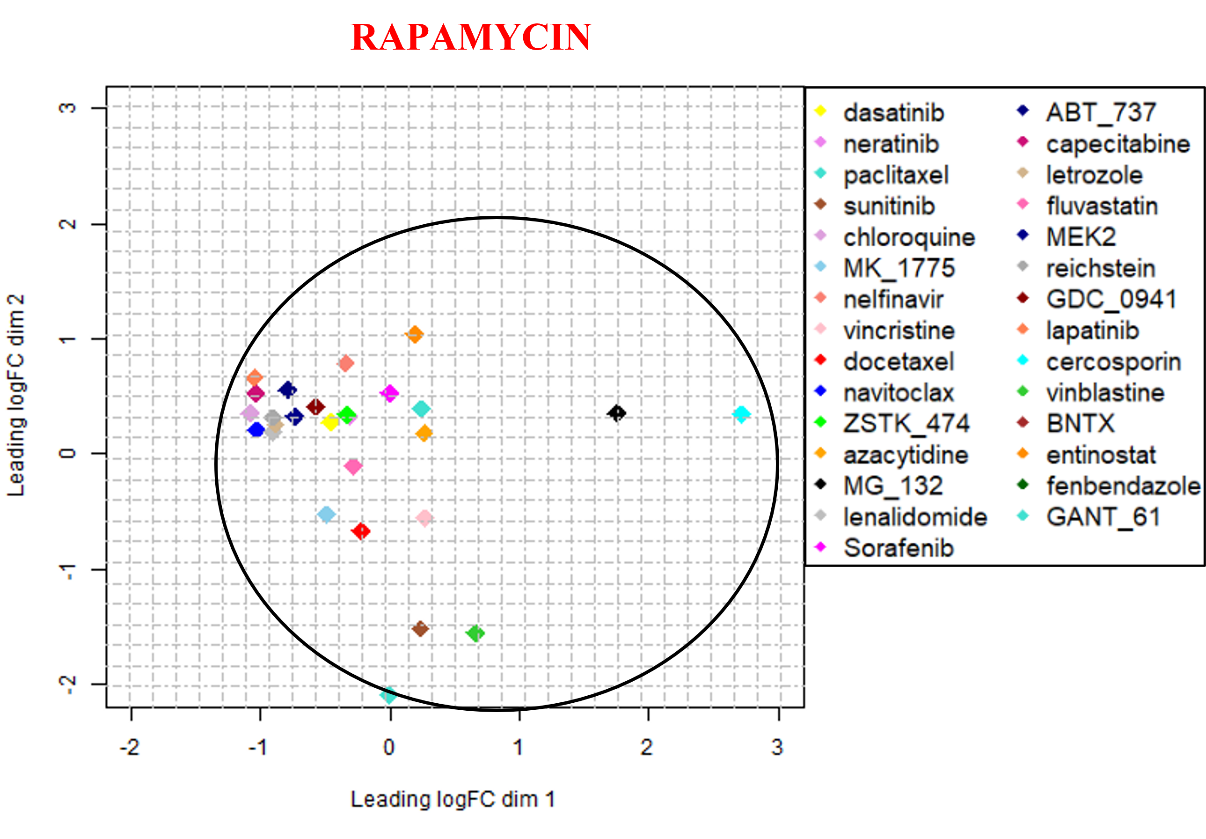

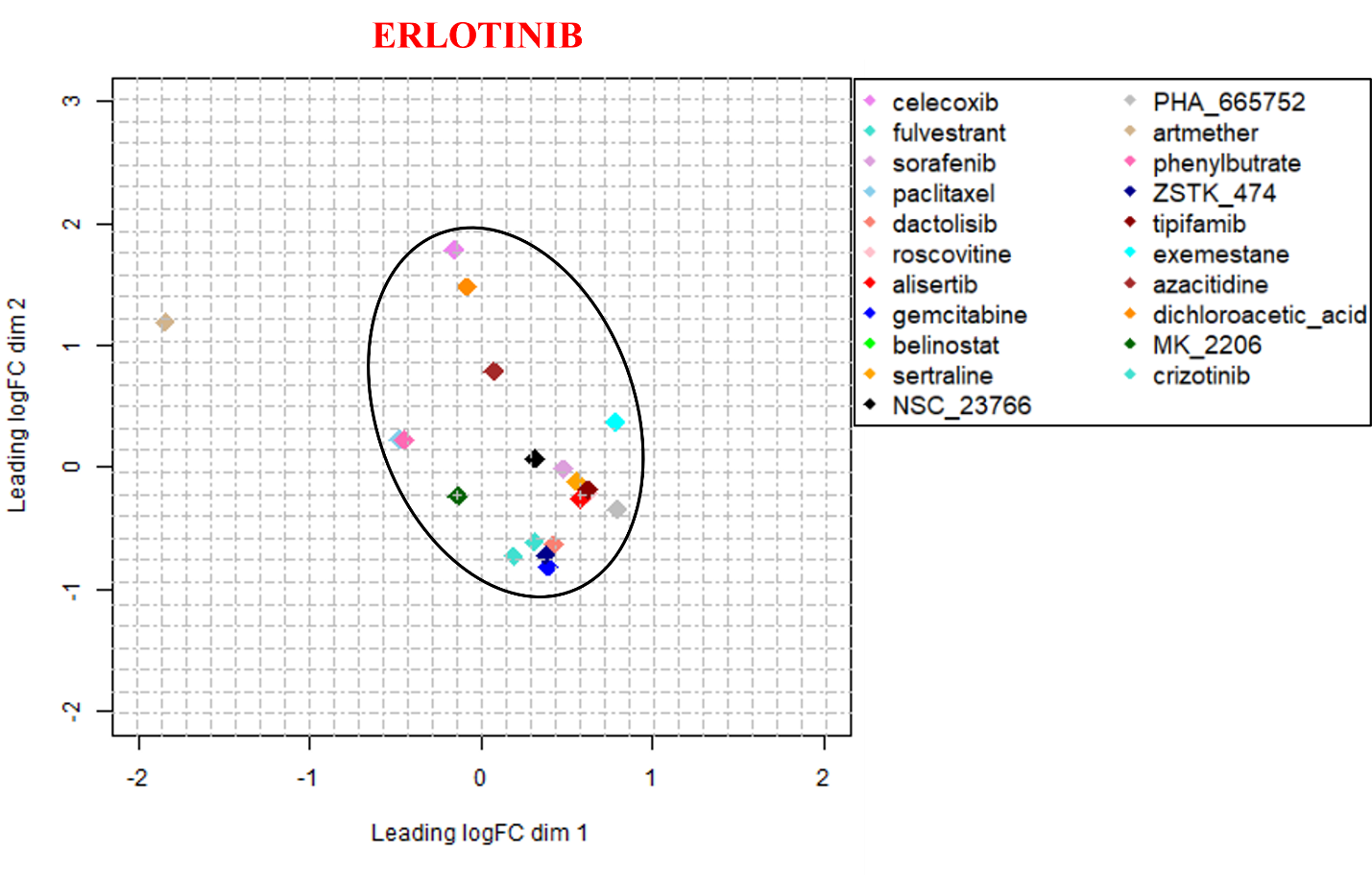

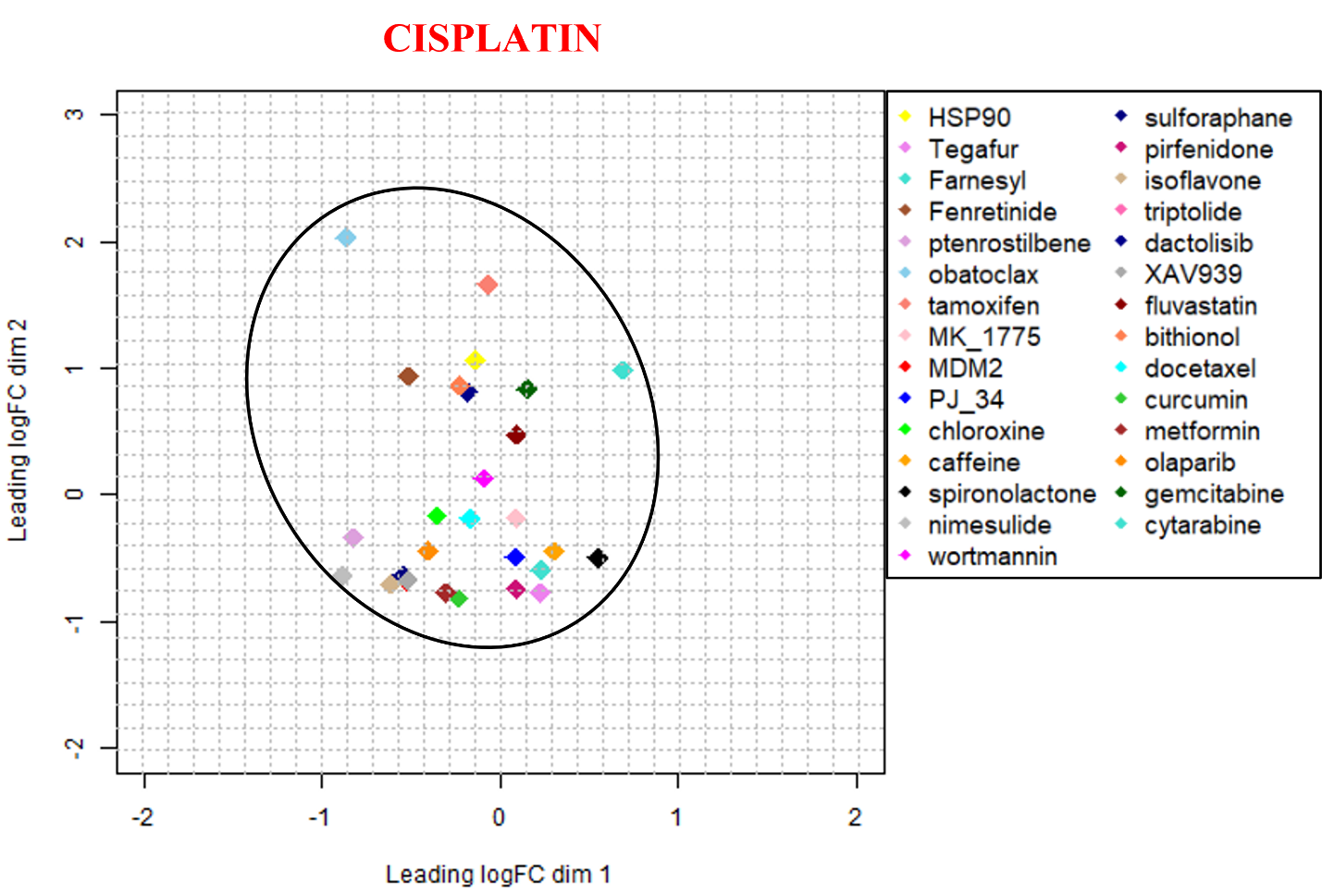

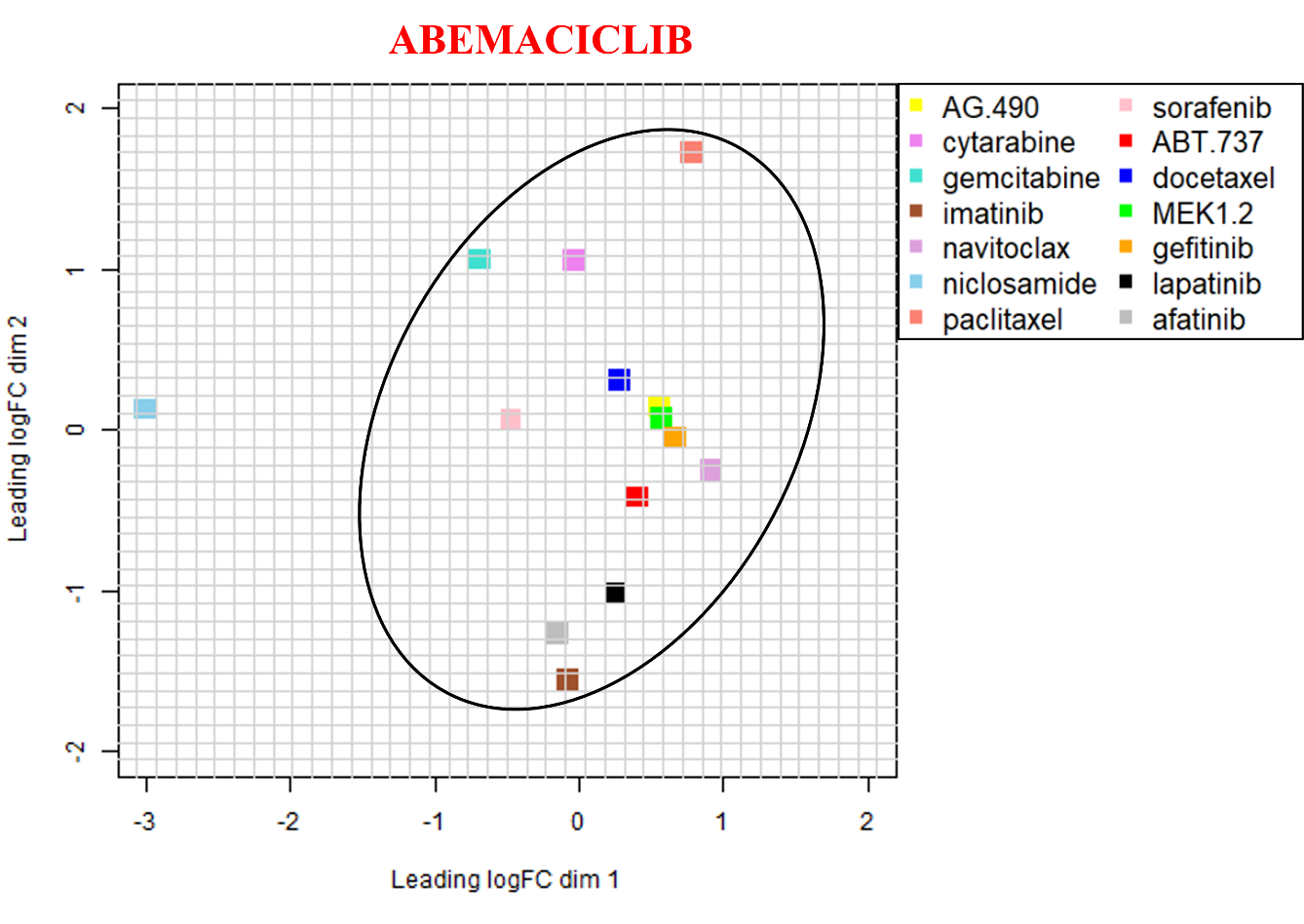

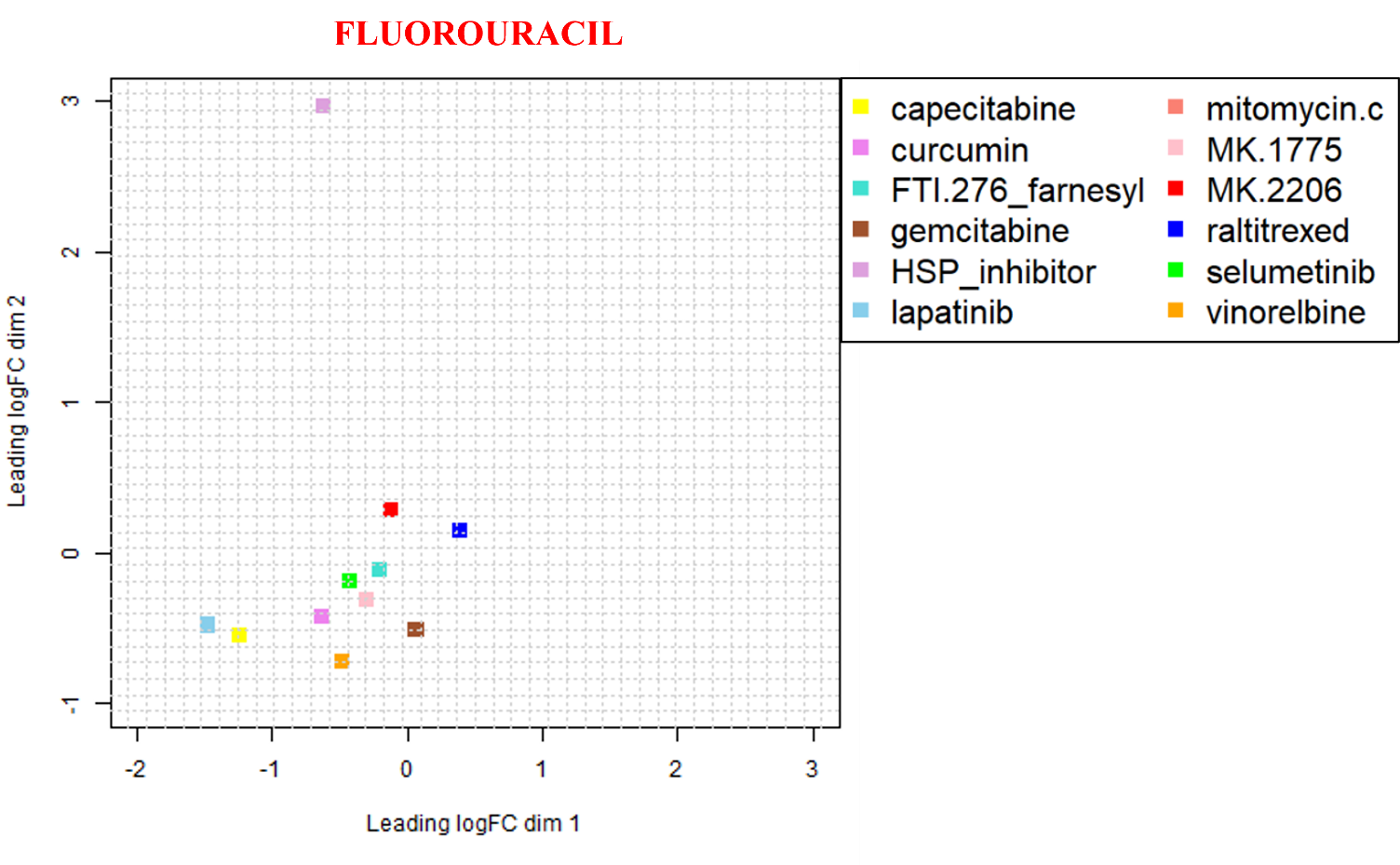
