## Supplementary figures and images for "Cancer Cell’s Seven Achilles Heels: Considerations for design of anti-cancer drug combinations"

### Figure_1A.png

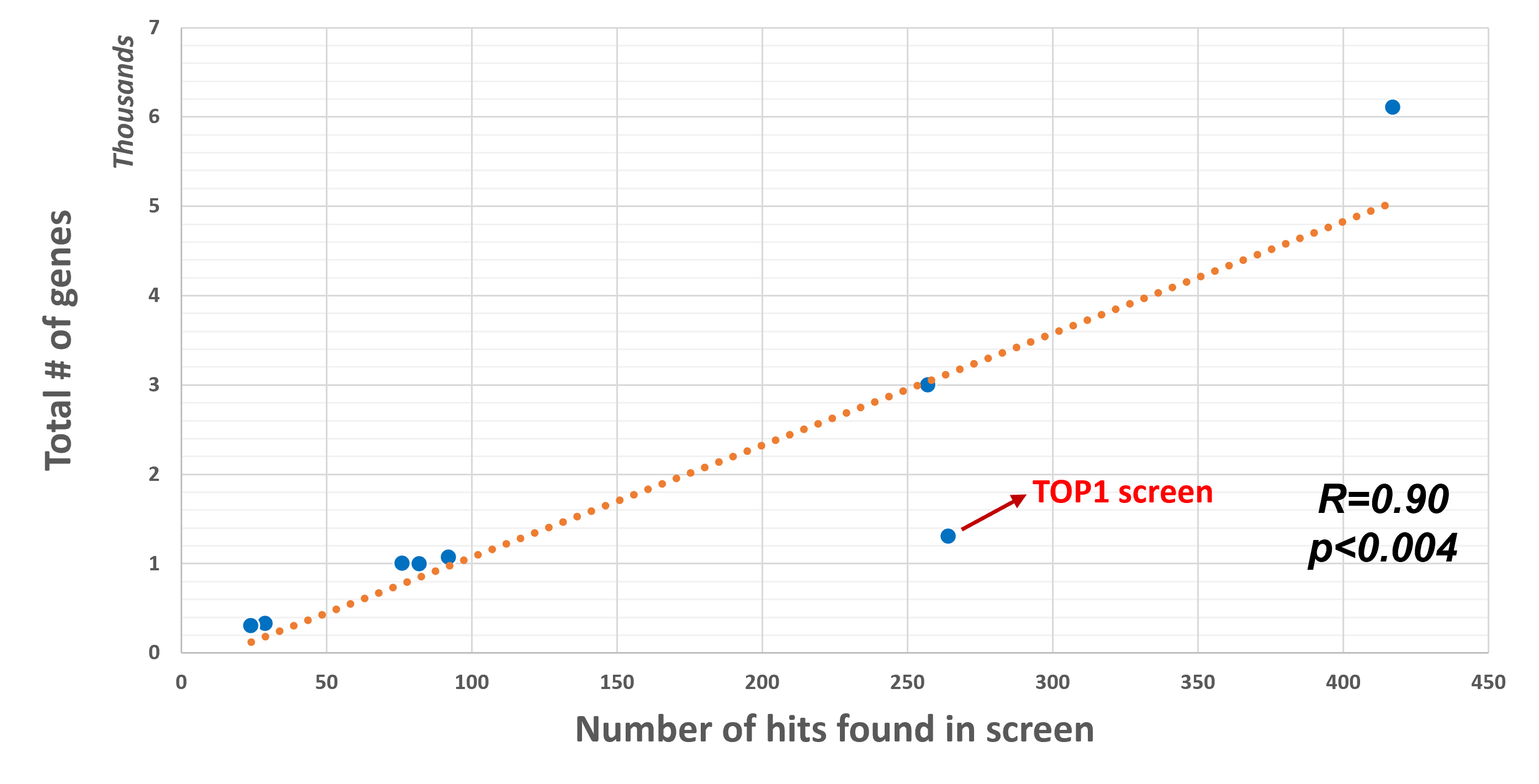

### Figure_1B.png

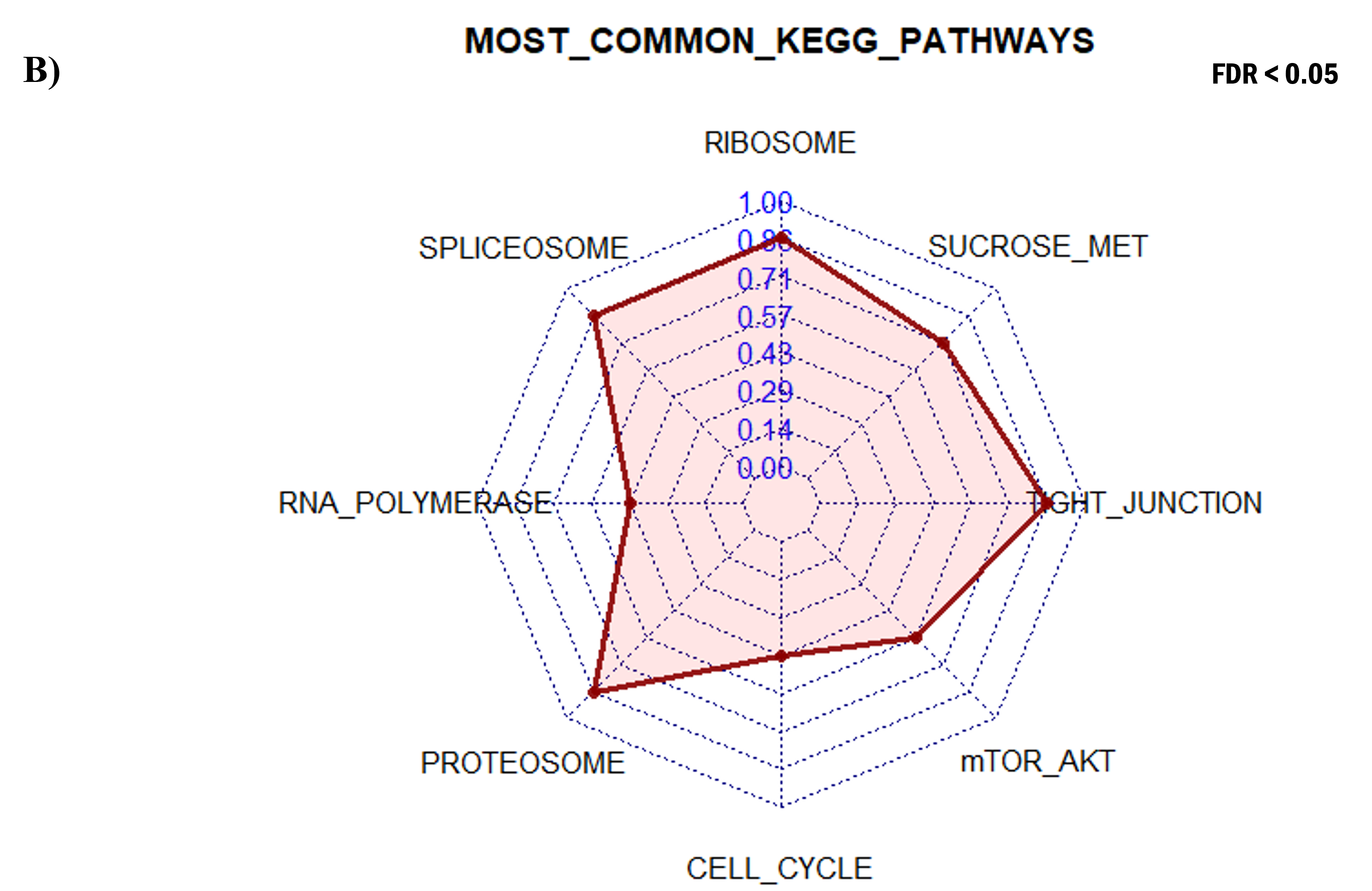

### Figure_1C.png

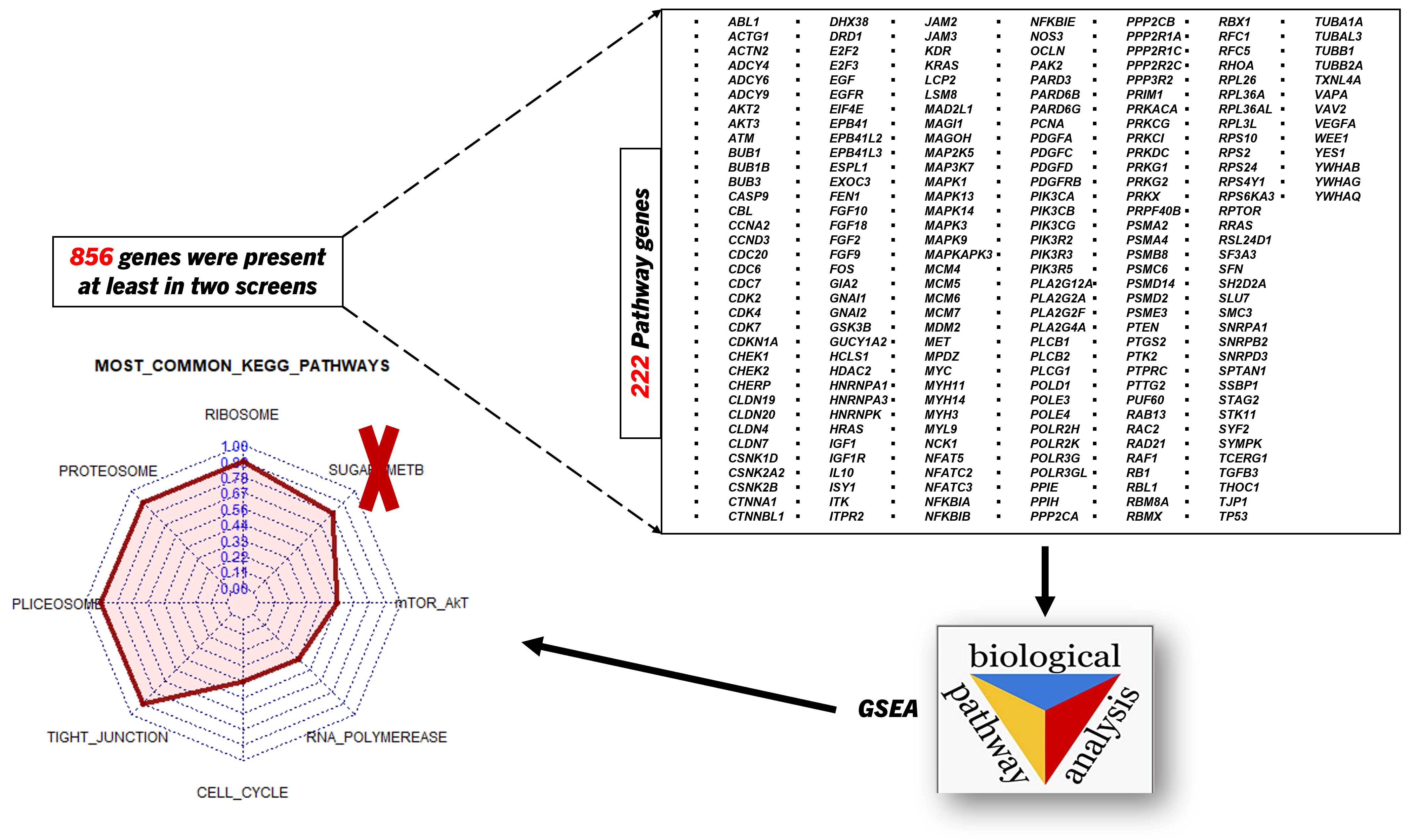

### Figure_2A.png

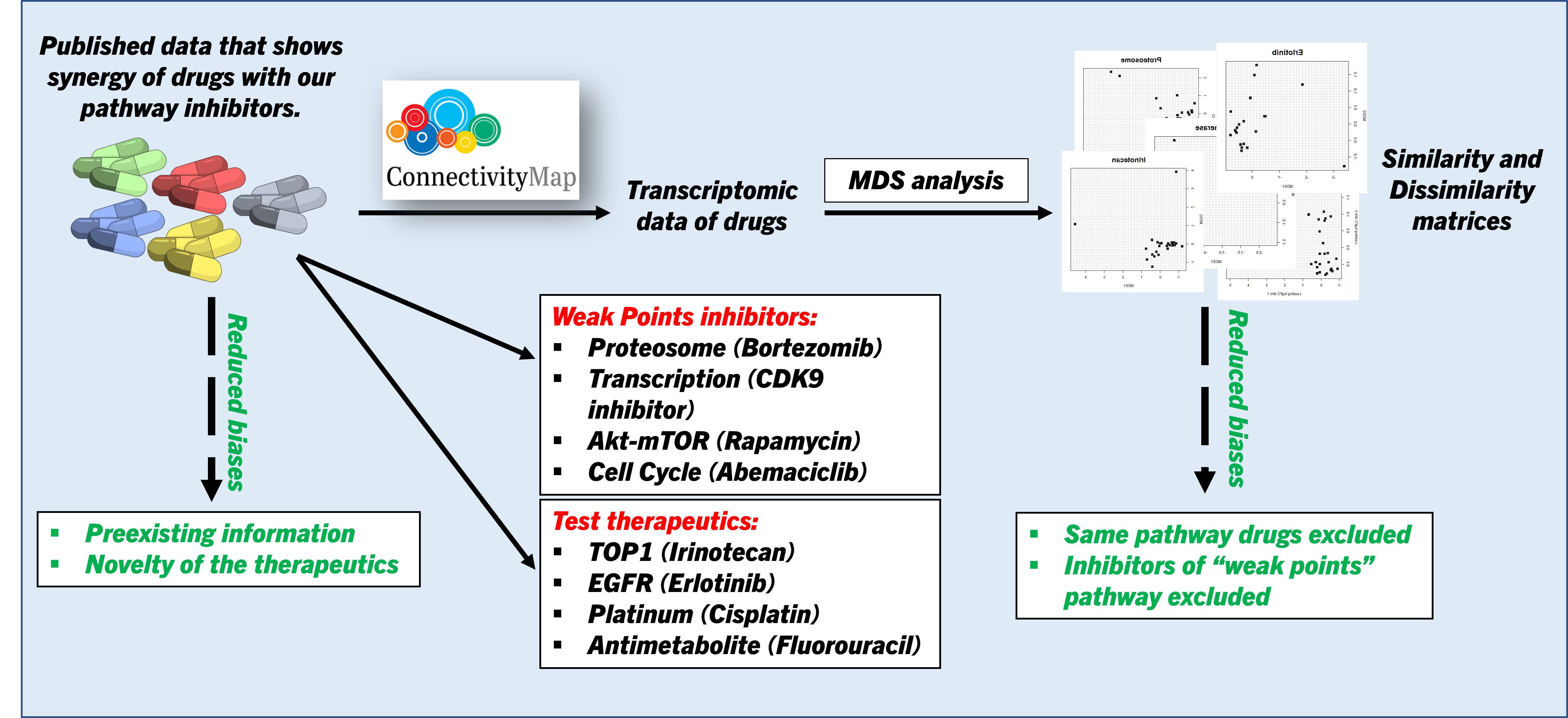

### Figure_2B_1.png

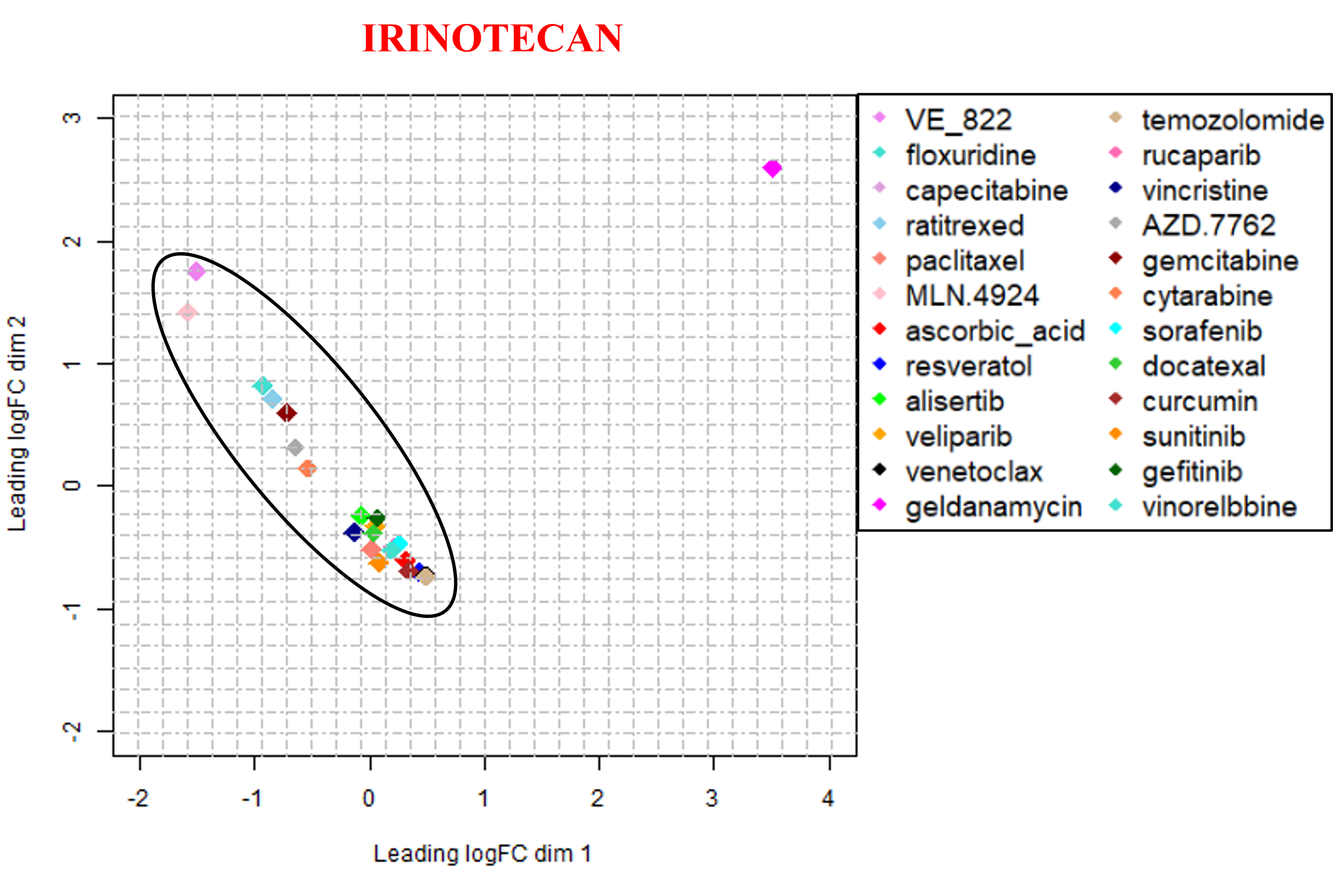

### Figure_2B_2.png

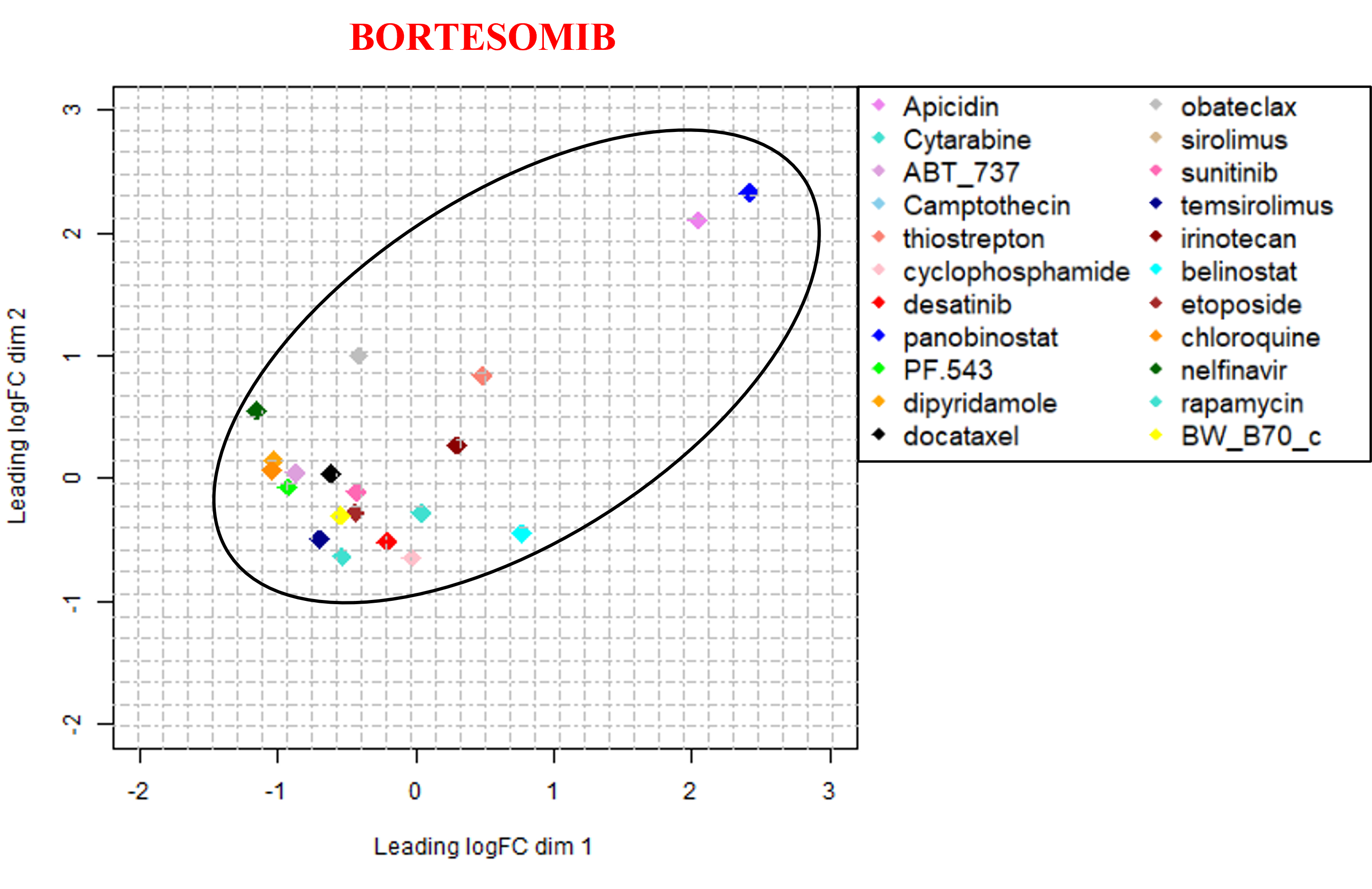

### Figure_2C.png

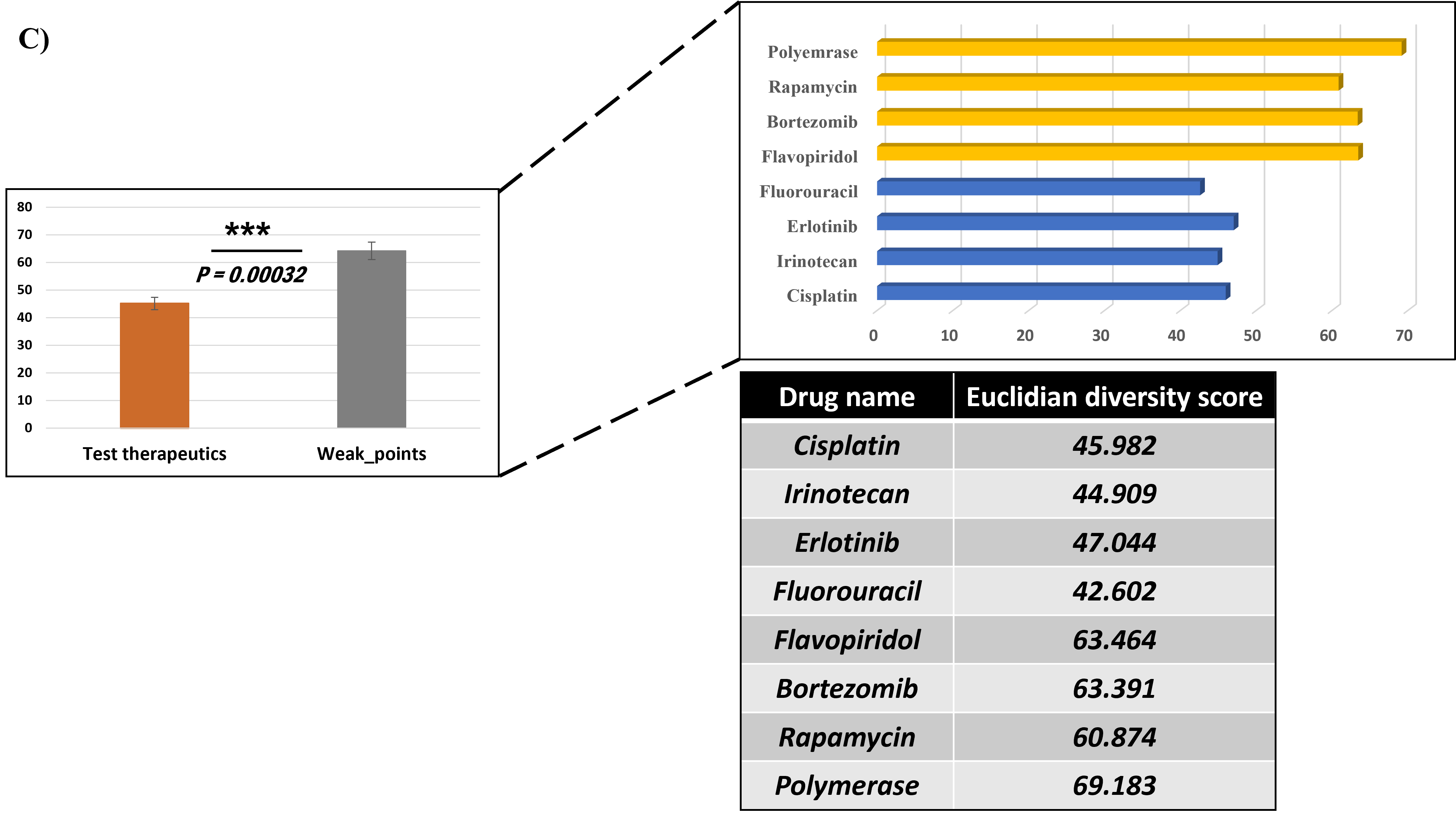
